# Visuospatial coding by theta oscillations in human hippocampus

**DOI:** 10.64898/2026.05.19.725196

**Authors:** Kenneth Rostowsky, Naoum P. Issa, Shasha Wu, James X. Tao, Hiba A. Haider, Sandra L. Rose, Peter C. Warnke, David Satzer, Rodrigo M. Braga, Stephan U. Schuele, Anna Shinn, Lingxiao Shi, Joel L. Voss, James E. Kragel

## Abstract

The hippocampus has been proposed to support visual processing and perception, challenging longstanding accounts that emphasize navigation or declarative memory. A key prediction of visual-processing accounts is that the hippocampus should exhibit similar visuospatial coding properties to those of higher-order visual neocortical areas, such as sensitivity to the size of visual stimuli and contralateral visual field biases. We tested for these properties using intracranial EEG to measure hippocampal neural activity during a retinotopic mapping task. The hippocampus exhibited characteristic slow (∼2 Hz) and fast (∼8 Hz) theta oscillations throughout the task. Fast theta was responsive to the presence but not the amount of visual stimulation. In contrast, slow theta did not generally respond to stimulus presence but scaled with the size of the visual stimulus, consistent with larger receptive fields. Slow theta also showed a contralateral bias, an effect that was specific to the right hippocampus. None of these effects were attributable to microsaccades or performance of the concurrent vigilance task. These findings provide electrophysiological evidence for visual field coding by human hippocampus, supporting accounts of hippocampal function that emphasize its role atop the visual hierarchy. Visual processing of this kind may combine with self-motion, memory, and other signals to support the broader spatial and mnemonic functions with which hippocampal theta oscillations have long been associated.

## Introduction

Functional accounts of the hippocampus and adjacent structures of the medial temporal lobes predominantly emphasize navigation [1, 2], as well as declarative [3, 4] and episodic memory [5–7]. However, some theories instead highlight its role in visual perception, owing to the position of the hippocampus as the apex of visual processing streams [8–12]. If the hippocampus is crucial for visual processing, as suggested by prior work using naturalistic stimuli [9, 13–16], one would expect it to respond to even basic visual stimulation in ways that reflect the organizational properties of other visual areas.

Visuospatial organization in the brain commonly follows two forms [17]. Retinotopic maps contain precise representations of the contralateral visual field and over-represent the field center [18]. Such maps are found in widespread visual regions [19, 20], including along the ventral visual stream [21]. Visual field biases, by contrast, are a form of visuospatial coding where neuronal populations are broadly tuned to certain locations [22], without clear map-like organization [17]. Recent functional MRI (fMRI) studies have suggested that the hippocampus may exhibit the latter form of organization, showing contralateral visual field biases [23–25] and topographic connectivity with visual cortex [26]. Whether these fMRI signals reflect direct visual drive to the hippocampus remains an open question, as the hemodynamic responses measured by fMRI are an indirect proxy for underlying neural activity [27–30].

Electrophysiological signals of hippocampal neural activity demonstrate visuospatial sensitivity in rodents [31] and nonhuman primates [32–34], but this relationship has not been tested in humans. The human hippocampus exhibits prominent theta-band oscillations in the local field potential, with distinct slow (∼2-5 Hz) and fast (∼5-10 Hz) theta thought to support distinct functions [35, 36]. Distinct regions along the hippocampal long axis support different functions [37, 38] and have different visuospatial tuning [39], with a posterior-anterior distinction following relative connectivity with parahippocampal cortex that conveys spatial and contextual information to posterior hippocampus [40–42] versus perirhinal cortex that conveys object and feature information to anterior hippocampus. Notably, fast theta encroaches upon the alpha band (∼8-12 Hz) in visual neocortex, where retinotopic organization is marked by contralateral alpha desynchronization and co-occurring increases in broadband high-frequency activity (HFA) [43]. Testing whether slow and fast hippocampal theta oscillations differ in their visuospatial sensitivity may thus help specify which hippocampal circuits are engaged by visual input, and whether the organizational properties of those circuits align with perceptual versus navigational or mnemonic functions.

We used intracranial EEG (iEEG) in human neurosurgical patients to characterize hippocampal responses during a retinotopic mapping task [20]. Our goals were to determine whether hippocampal oscillations were driven by visual input, and if so, whether their spatial sensitivity was consistent with visual field coding. Because the hippocampus is sensitive to attention and task engagement [44, 45], we assessed microsaccades that indicate lapses in engagement and shifts in the visual field [46, 47] and behavioral performance on the concurrent change-detection task as potential alternative explanations for changes in hippocampal activity. Our experiment thus provides a strong test of whether human hippocampal electrophysiology shows evidence for visuospatial organization, with implications for perceptual accounts of hippocampal function.

## Results

### Task performance during retinotopic mapping

Participants (n = 12) performed a retinotopic mapping task [20] adapted for use with iEEG. Simple shapes such as wedges, bars, and annuli comprised of an animated texture of objects swept across the visual field in various motion paths. Participants were instructed to maintain visual fixation on a centrally located dot that changed color every 1-5 s and to indicate color changes via button-press responses (Fig. 1A). Participants responded to 86*±*3% (mean ± SEM) of color changes with a median response time of 447±36 ms (Fig. 1B-C). Simultaneous eye-movement tracking indicated that 89*±*3% of fixations were within 2 degrees of visual angle of the central dot (Fig. 1D). These findings indicate that participants maintained central fixation and vigilance throughout most of the task, providing a basis to examine visuospatial organization of hippocampal activity.

**Figure 1:**
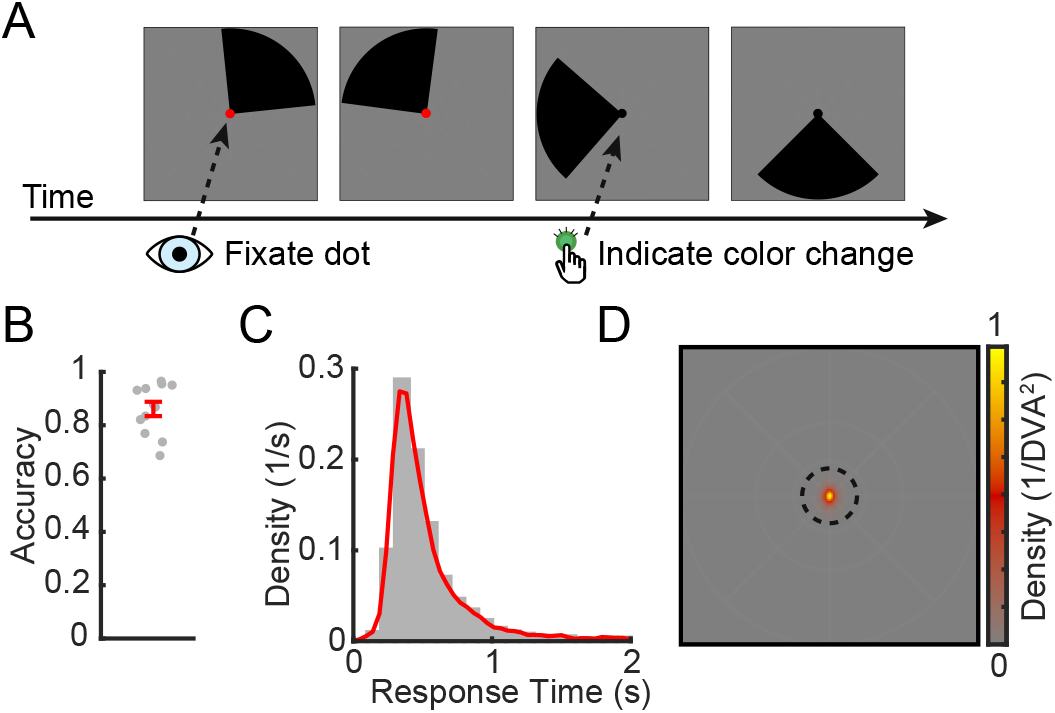
Retinotopic mapping task. (A) Example images from a trial in which a wedge-shaped mapping stimulus swept around the screen while participants maintained visual fixation on a central dot and indicated with a button-push response when the dot changed color. (B) Mean accuracy per participant (gray dots) and across the group (red error bars indicate SEM). (C) Response-time distribution for all participants and trials combined. Red line indicates a kernel density estimate. (D) Heatmap indicates the average distribution of gaze throughout the task, measured per squared degree of visual angle (DVA). Dashed circle indicates 2 DVA around the central dot, which contained the majority of all fixations.

### Hemispheric asymmetry of slow and fast hippocampal theta oscillations

We used a detection algorithm [48, 49] to identify neural oscillations during task performance, focusing on slow and fast theta oscillations based on their prevalence in human hippocampal iEEG [36] (see Fig. 2A for representative signals from a single participant). To summarize oscillatory activity across all 108 hippocampal contacts, we applied a data-driven clustering algorithm (see Methods and Fig. S1 for details) to identify channels with similar oscillatory activity to one another. Oscillatory activity was well described by two clusters of contacts: one primarily exhibiting slow theta oscillations near 2 Hz and the other exhibiting fast theta oscillations near 8 Hz (Fig. 2B).

**Figure 2:**
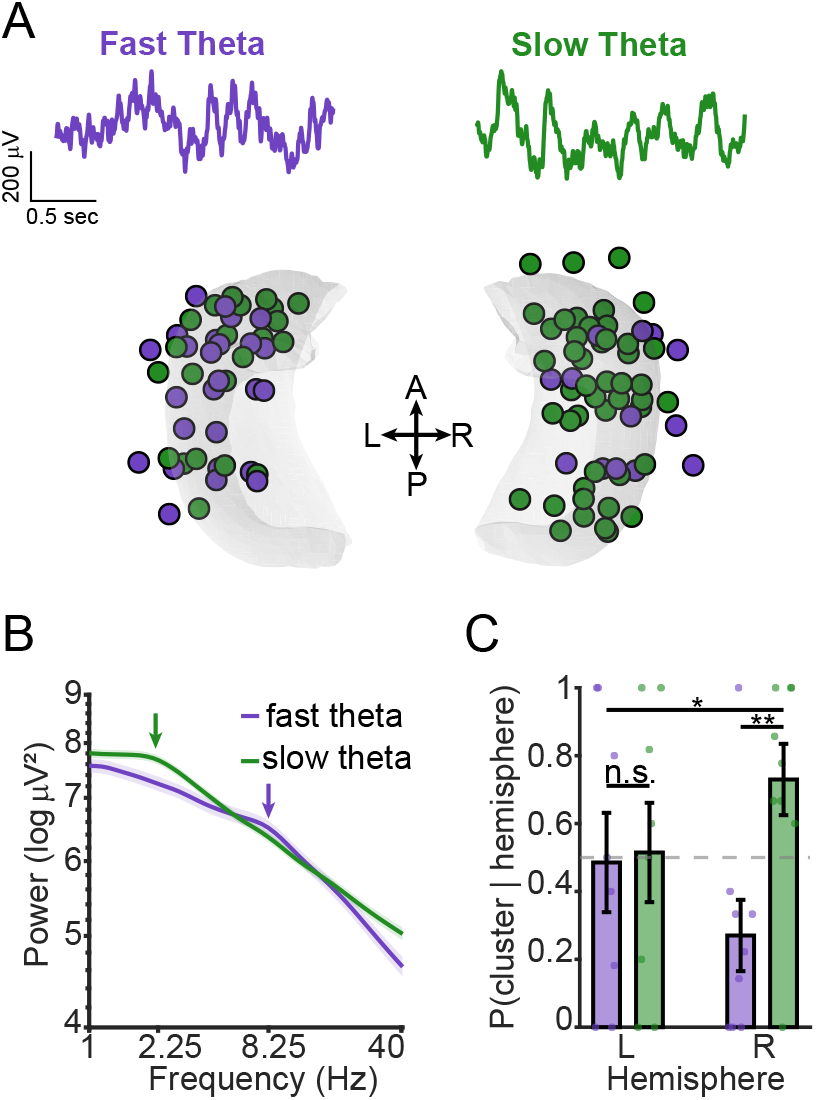
Distinct hippocampal slow and fast theta oscillations during visual-field mapping. (A) Above, example hippocampal traces, showing fast (purple) and slow (green) theta during task performance. Below, hippocampal contacts colored based on oscillatory activity (*N* = 108 channels; 39 with fast theta and 69 with slow theta). (B) Group-averaged power spectra from two clusters of channels showing prominent slow and fast theta oscillations, shaded bands indicate bootstrap-estimated SEM across subjects. Arrows indicate locations of corresponding peak frequencies in slow and fast theta. (C) Hemispheric differences in slow and fast theta. Contacts with slow theta were more prevalent in the right hemisphere. Dashed line indicates expected distribution of contacts in each cluster per hemisphere. Error bars denote bootstrap-estimated SEM. *, *p <* 0.05, ** *p <* 0.005.

Slow and fast theta contacts were distributed asymmetrically across hemispheres (Fig. 2C). Within the left hippocampus, contacts were equally likely to exhibit slow or fast theta (54 *±* 5% vs. 46 *±* 5%; odds = 1.2, *p* = 0.66, permutation test), whereas the right hippocampus favored slow theta (78 *±* 4% vs. 22 *±* 4%; odds = 3.6, *p* < 0.0001, permutation test). Accordingly, right hippocampal contacts were roughly three times more likely to exhibit slow theta than left hippocampal contacts (odds ratio = 3.03, *p* = 0.045, permutation test). The slow and fast theta clusters did not differ significantly in their anterior-posterior distribution (rank-biserial difference = −0.24, permutation *p* = 0.15), suggesting comparable spatial spread along the long axis of the hippocampus regardless of cluster membership (see Fig. S1D-E for additional details). Together, these findings replicate the presence of distinct slow and fast theta oscillations in the human hippocampus, with an asymmetry favoring slow theta in the right hemisphere. We next asked whether these oscillatory regimes differ functionally in their sensitivity to visual stimulation.

### Fast and slow theta oscillations show distinct responses to visual stimulation

We first asked how theta oscillations in the hippocampus responded to visual stimulation in the task. We compared theta power observed during periods with the retinotopic mapping stimulus (ON) to periods when subjects were similarly engaged in the task, but without the retinotopic mapping stimulus (OFF, Fig. 3A). Because hippocampal responses may be spatially specific and vary by participant, analyses were first performed for each contact independently. Slow theta was sensitive to ON versus OFF periods in 13% of hippocampal contacts, with a mixture of increases and decreases (Fig. 3B).

**Figure 3:**
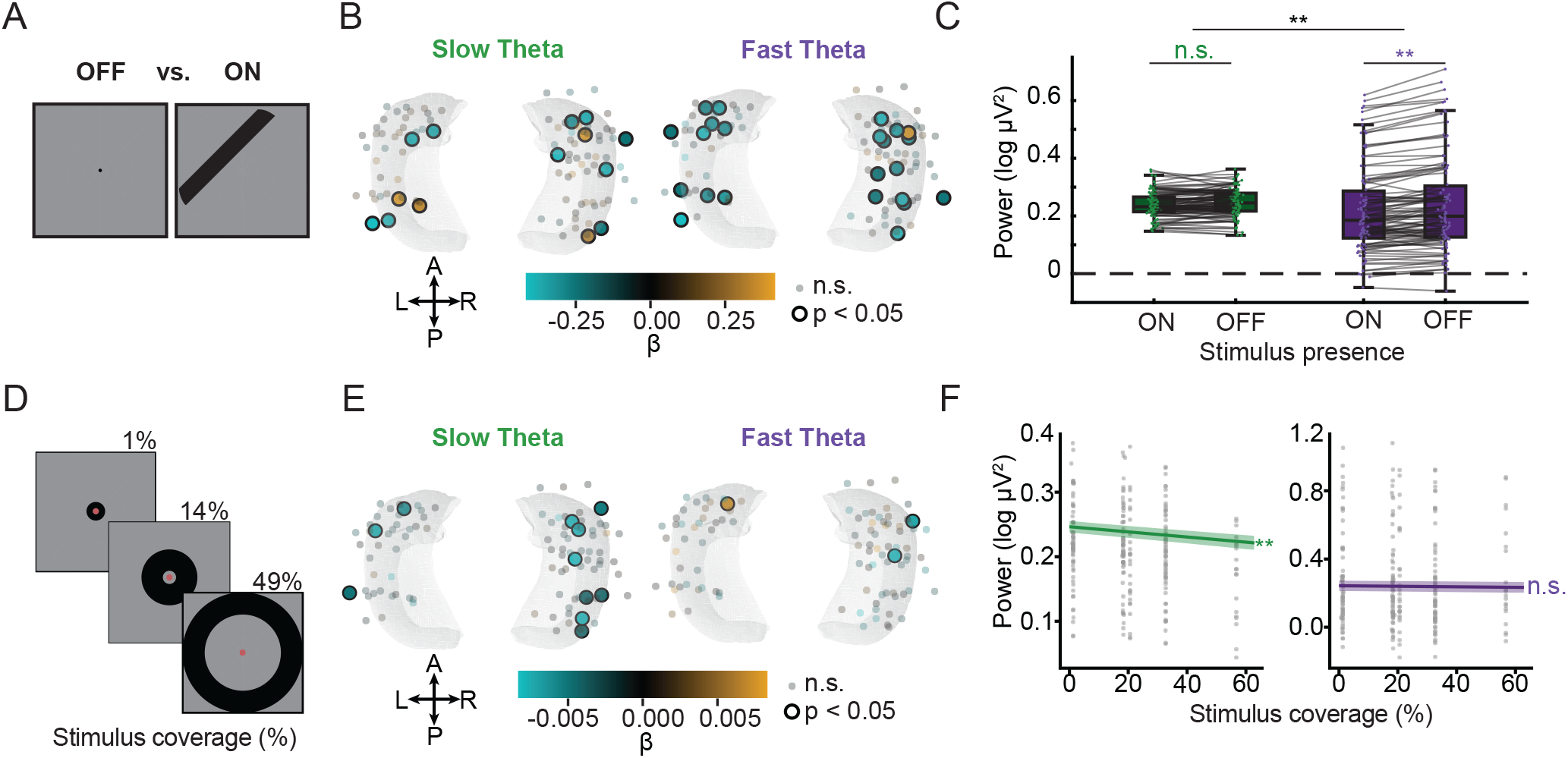
Slow and fast theta have distinct responses to visual stimulation. (A) Schematic overview of comparison between ON and OFF periods. ON periods occur when any retinotopic mapping stimuli are present in the visual field, OFF periods occur in the absence of retinotopic mapping stimuli. (B) Contact-level comparisons of mean oscillatory power between ON and OFF periods for slow theta (left) and fast theta (right). Contacts with a significant effect are indicated by a larger circle with a thick black border. (C) Group-level comparison of mean oscillatory power between ON and OFF periods for slow theta (left, green) and fast theta (right, purple). Each gray line indicates measurements taken from the same hippocampal contact. (D) Schematic overview of the stimulus coverage present during each trial, reflecting varying amounts of the central display occupied by retinotopic mapping stimuli. (E) Contact-level results showing standardized beta values for slow theta (left) and fast theta (right). (F) Group-level relationship between percent visual stimulation and oscillatory power for slow theta (left, green) and fast theta (right, purple). Lines indicate fixed effects from LMEMs; shaded regions indicate SEM. *, *p <* 0.05, **, *p <* 0.005, Holm-Bonferroni corrected, parametric bootstrap.

Group-level analyses across all participants and hippocampal contacts used linear mixed effects modeling (LMEM), with separate models fit for slow and fast theta. These models examined how visual stimulus presence (whether any stimulus was on the screen), coverage (proportion of the screen occupied by a visual stimulus), and laterality (the difference in contralateral vs. ipsilateral coverage by the visual stimulus) each uniquely related to theta power. LMEM showed no significant relationship between slow theta power and stimulation presence (*χ*^2^(1) = 0.99, *p* = 1.00, Holm-Bonferroni corrected, parametric bootstrap), consistent with our contact-level analysis showing no consistent ON/OFF response pattern.

In contrast, visual stimulation consistently affected fast hippocampal theta, with stimulus presence significantly influencing 21% of contacts (Fig. 3B). All but one significant contact showed decreases in fast theta during ON versus OFF periods, and LMEM confirmed significant decreases at the group level (*χ*^2^(1) = 96.6, *p* = 0.0028, Holm-Bonferroni corrected, parametric bootstrap). To directly quantify the qualitative difference between fast and slow theta, we fitted another model with frequency as an interaction term. We found that ON/OFF sensitivity was significantly greater for fast compared to slow theta (*χ*^2^(1) = 33.8, *p <* 0.001, parametric bootstrap). We did not find evidence that the hemisphere of the hippocampal recording impacted ON/OFF sensitivity in general (*χ*^2^(1) = 0.04, *p* = 0.84, parametric bootstrap) or with regards to frequency differences (*χ*^2^(1) = 1.21, *p* = 0.27, parametric bootstrap). Together, these findings show that sensitivity to overall stimulus presence is specific to fast theta oscillations.

Having established that fast theta selectively responds to the presence of visual stimulation, we next asked whether slow or fast theta were sensitive to the extent of stimulation. Sensitivity to stimulus size would be expected to differ depending on the underlying spatial tuning. Signals reflecting coarse or spatially diffuse representations (e.g., larger effective receptive fields) should scale with the amount of visual input, whereas signals primarily responding to the presence or onset of stimulation may not.

To test this, we quantified stimulus coverage as the proportion of the screen occupied by the visual stimulus during each trial (Fig. 3D). We found that slow theta correlated negatively with stimulus coverage (i.e., lower slow theta associated with greater coverage). At the contact level, 11% of contacts showed diminished slow theta power with increasing stimulus coverage (Fig. 3E). For fast theta, only 3% of contacts were significantly modulated by coverage. At the group level (Fig. 3F), the LMEM described above identified a significant effect of stimulus coverage for slow theta (*χ*^2^(1) = 62.9, *p* = 0.0028, Holm-Bonferroni corrected, parametric bootstrap) but not fast theta (*χ*^2^(1) = 3.85, *p* = 0.50, Holm-Bonferroni corrected, parametric bootstrap). Testing for differences between slow and fast theta, we found that this sensitivity was greater for slow theta (*χ*^2^(1) = 7.07, *p* = 0.008, parametric bootstrap).

Taken together, the aforementioned results indicate functional dissociation of slow versus fast theta bands. Fast 4 theta was generally sensitive to whether visual stimulation was present, whereas slow theta was sensitive to the magnitude of visual input.

### Hippocampal theta oscillations show a contralateral visual field bias

To test whether hippocampal theta has contralateral biases to visual stimulation, we assessed whether laterality in visual stimulation modulated theta oscillations (Fig. 4A). Contralateral visual hemifield stimulation affected slow theta oscillations. There were significant decreases in slow theta power during contralateral visual stimulation in 6% of contacts, with more located in the right than left hemisphere (Fig. 4B). Group-level analyses using the same LMEM described above indicated no overall relationship of slow theta power to visual stimulus laterality (*χ*^2^(1) = 4.20, *p* = 0.37, Holm-Bonferroni corrected, parametric bootstrap), but significant interaction with hemisphere, such that the correlation of slow theta power with stimulus laterality was significantly different for right versus left hippocampus (*χ*^2^(1) = 14.6, *p* = 0.0028, Holm-Bonferroni corrected, parametric bootstrap). This reflected a significant effect of stimulus laterality on slow-theta power in right hippocampus (*β* = −1.90*e*-4, *SE* = 4.1*e*-5, *χ*^2^(1) = 21.05, *p* = 0.0024, Holm-Bonferroni corrected, parametric bootstrap) but not left hippocampus (*β* = 5.8*e* - 5, *SE* = 5.0*e*-5, *χ*^2^(1) = 1.3, *p* = 0.77, Holm-Bonferroni corrected, parametric bootstrap).

**Figure 4:**
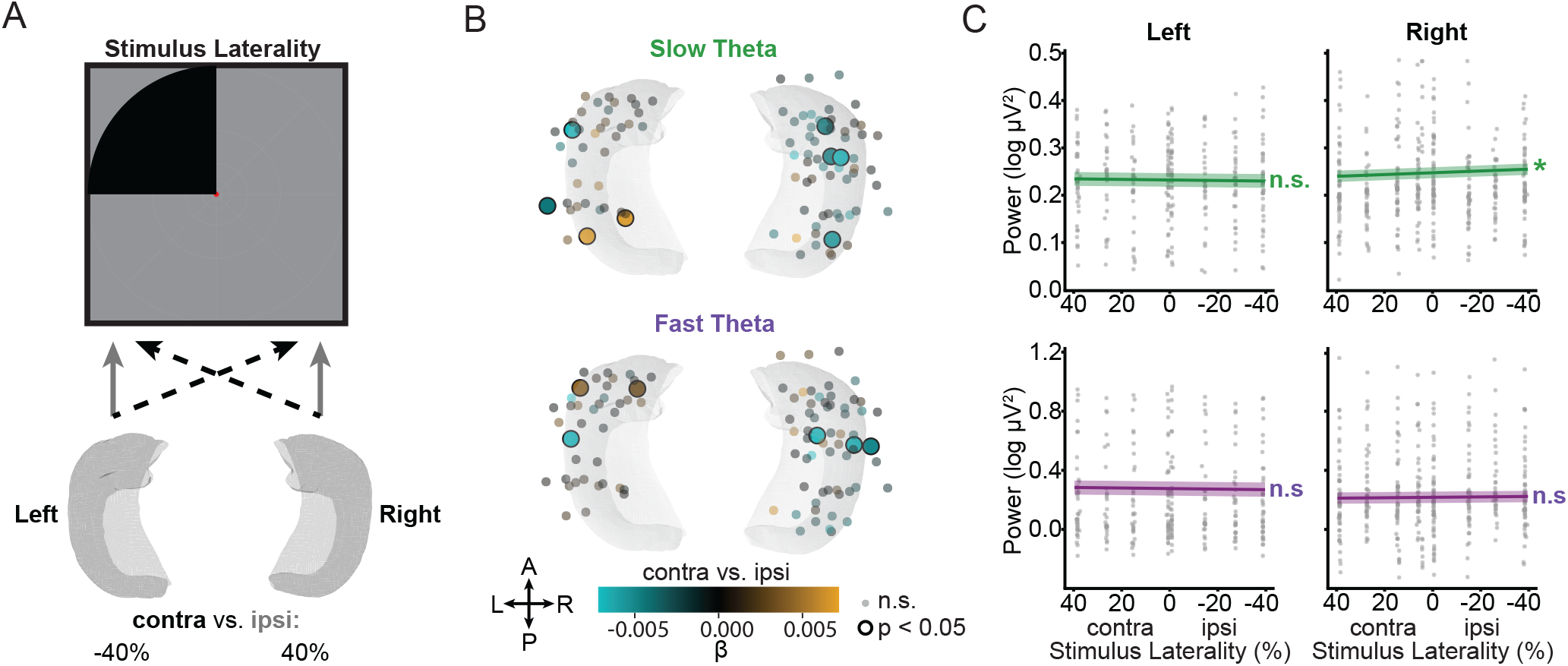
Slow theta oscillations show hemifield organization. (A) Schematic overview of hypothesized relationship between theta power and contralateral visual hemifield stimulation. Greater contralateral visual hemifield stimulation is hypothesized to decrease theta power. (B) Contact-level results showing standardized beta values for slow (green, top) and fast (purple, bottom) theta power as a function of percent contralateral visual hemifield stimulation. Contacts where theta power is significantly related to laterality in visual hemifield stimulation are indicated by a larger circle with a thick black border. (C) Group-level relationship for slow (green, top) and fast (purple, bottom) theta power as a function of percent contralateral visual hemifield stimulation, stratified by hippocampal contact hemisphere. Negative stimulus laterality indicates percent of stimulus in the ipsilateral visual hemifield for a given hippocampal contact. Gray dots represent individual observations for each hippocampal contact. Dashed line indicates equal visual hemifield stimulation or lack thereof. Solid lines indicate fixed effects from LMEMs; shaded regions indicate SEM. *, *p <* 0.05, Holm-Bonferroni corrected, parametric bootstrap.

Fast theta power showed a similar, albeit weaker, contralateral bias, with 4% of contacts showing contralateral decreases in relation to laterality (Fig. 4B). At the group level (Fig. 4C), LMEM indicated no significant differences in the relationship of fast theta with stimulus laterality (*χ*^2^(1) = 0.2248, *p* = 1.00, Holm-Bonferroni corrected, parametric bootstrap). As was the case for slow theta, there was a significant relationship between fast theta power and stimulus laterality that varied significantly by hemisphere (*χ*^2^(1) = 13.3, *p* = 0.0088, Holm-Bonferroni corrected, parametric bootstrap). Stimulus laterality had opposing effects on fast theta power depending on hippocampal hemisphere, however, neither left nor right hippocampal fast theta power alone had a significant relationship with stimulus laterality. In the right hippocampus, there were decreases in fast theta with increasing contralateral stimulation (*β* = −1.4*e*-4, *SE* = 5.9*e*-5, *χ*^2^(1) = 6.09, *p* = 0.073, Holm-Bonferroni corrected, parametric bootstrap) but in the left hippocampus, there were decreases in fast theta with increasing ipsilateral stimulation (*β* = 1.9*e*-4, *SE* = 7.0*e*-5, *χ*^2^(1) = 7.21, *p* = 0.052, Holm-Bonferroni corrected, parametric bootstrap).

When comparing between fast and slow theta oscillations, there was no significant difference in the effect of visual laterality (*χ*^2^(1) = 0.0356, *p* = 0.85, parametric bootstrap). However, the contralateral bias varied significantly for left versus right hippocampus (*χ*^2^(1) = 19.39, *p <* 0.001, parametric bootstrap), without significant variation by the frequency of theta (*χ*^2^(1) = 1.21, *p* = 0.27, parametric bootstrap). These findings indicate that both slow and fast theta had similar visual-field bias that differed by hemisphere, with contralateral bias for right hippocampus.

### Hippocampal theta is not predicted by task engagement or fixational eye movements

The observed contralateral biases in hippocampal theta and visual sensitivity in fast theta could potentially reflect the influences of ongoing task-related behaviors. Specifically, if central attention or anticipation of an upcoming color change modulates theta, our findings could reflect task engagement rather than visual responses. Hippocampal theta and HFA are known to be sensitive to attentional state and stimulus expectancy [44, 45]. Saccadic eye movements are also known to modulate ongoing theta rhythms [49–52]. This is particularly relevant as the rate of microsaccades is known to vary with the expectation of upcoming, task-relevant stimuli [53]. To test whether theta is sensitive to such factors during our task, we examined slow and fast theta power as a function of two trial-level variables: whether microsaccades were present during a trial and whether participants made accurate responses.

We quantified microsaccades using a well-established algorithm [54, 55]. Participants made microsaccades at a rate of 1.89 ± 0.34 Hz, consistent with prior work [56, 57]. We did not find evidence that these fixational eye movements were directed toward the center of the stimulus (group mean cos*θ* = 0.038, permutation null 95% CI [0.028, 0.041], *p* = 0.13 two-tailed) or the central fixation dot (group mean cos*θ* = -0.020, permutation null 95% CI [-0.023, 0.029], *p* = 0.96 two-tailed). We also confirmed that fixational eye movements were not generally aligned toward or away from the center of the aperture towards the central fixation dot (mean |cos*θ*| = 0.64, 95% CI [0.64, 0.65], *p* = 0.47). As such, we focused our analyses on the execution of microsaccades in general, without focusing on directional effects.

We found no relationship between accurate responses and slow theta power (*χ*^2^(1) = 2.53, *p* = 0.499, Holm Bonferroni corrected, parametric bootstrap) or fast theta power (*χ*^2^(1) = 0.131, *p* = 1.000, Holm-Bonferroni corrected, parametric bootstrap). Microsaccades were similarly unrelated to slow theta (*χ*^2^(1) = 0.411, *p* = 1.000, Holm-Bonferroni corrected, parametric bootstrap) and fast theta (*χ*^2^(1) = 0.152, *p* = 1.000, Holm-Bonferroni corrected, parametric bootstrap) power. There were no interactions between response accuracy and microsaccades on slow theta (*χ*^2^(1) = 0.400, *p* = 1.000, Holm-Bonferroni corrected, parametric bootstrap) or fast theta (*χ*^2^(1) = 3.77, *p* = 0.407, Holm-Bonferroni corrected, parametric bootstrap) power. In sum, we found no evidence that theta modulations were explained by task performance or ongoing oculomotor behavior.

### No sensitivity of hippocampal HFA to visual stimulation

We subjected hippocampal HFA values to the same analyses as described above for slow and fast theta oscillations. No group-level LMEM terms were significant (all *p* values *>* 0.05, Holm-Bonferroni corrected, parametric bootstrap), indicating that HFA was not sensitive to the presence, magnitude, or laterality of visual stimuli in either left or right hippocampus.

### Population receptive field modeling

To test whether hippocampal signals exhibited visuospatial organization, we fitted a pRF model to slow theta, fast theta, and broadband HFA observed at individual contacts using the compressive spatial summation (CSS) framework [58]. Across all three signal types, pRF model fits yielded average *R*^2^ values near zero (slow theta: *R*^2^ = 0.038 *±* 0.043, fast theta: *R*^2^ = 0.029 *±* 0.034, HFA: *R*^2^ = 0.026 *±* 0.023 [mean *±* SEM]), indicating that none of these signals were well described by a Gaussian receptive field model.

## Discussion

We used iEEG to test whether human hippocampal theta oscillations carry visuospatially organized signals. Fast theta was desynchronized by visual stimulation but showed no sensitivity to stimulus extent. In contrast, slow theta was not sensitive generally to visual stimulation, but decreased in step with the amount of the visual field stimulated, suggesting a larger receptive field than fast theta. Both fast and slow theta had greater decreases for contralateral compared to ipsilateral stimulation, primarily in right-hemisphere hippocampus. These findings suggest that both fast and slow hippocampal theta were visually sensitive and had visual field bias, with differences in the size of stimuli required to drive theta desynchronizations. Neither fast nor slow theta were modulated by microsaccades or accuracy on the ongoing change-detection task. These findings provide direct electrophysiological evidence that human hippocampal theta carries a coarse visual field bias, and thus that hippocampus supports visual processing.

Our findings were unexpected in light of prevailing accounts of hippocampal theta function. Fast theta is typically associated with posterior hippocampus, navigation, and spatial coding, while slow theta is associated with anterior hippocampus and memory encoding [35, 36]. Our results instead indicated that both fast and slow theta were responsive to the visuospatial properties for non-mnemonic and non-navigational stimuli, with distinctions for left versus right, rather than anterior versus posterior, hippocampus. The nonspecific visual sensitivity of fast theta, which responds to stimulus presence but not to where or how much of the field was stimulated, may reflect nonspecific visual drive arriving from neocortical areas. The scaling of slow theta with stimulus coverage, by contrast, is more consistent with a system acting on information integrated across the visual field, consistent with the spatially diffuse receptive fields characteristic of representations near the apex of the ventral visual hierarchy [59, 60]. Because the stimuli used in our task contained images known to activate the ventral visual stream [61, 62], slow theta’s spatial sensitivity may be routed through perirhinal and entorhinal inputs that convey ventral stream object and scene information to anterior hippocampus [11, 12]. Whether this specificity to slow theta reflects a general property of hippocampal circuitry or a consequence of using stimuli known to drive the ventral stream remains an important open question. In either case, both slow and fast theta responded significantly more to contralateral than ipsilateral stimuli, suggesting similar visual field biases in both signals.

Alternatively, the observed decreases in theta may reflect suppression of hippocampal processes involved in spatial encoding during externally driven visual stimulation [35, 63]. Importantly, this suppression was spatially specific. Slow theta decreases scaled with stimulus extent and both fast and slow theta exhibited contralateral bias, indicating that the suppression itself preserved information about the spatial structure of visual input. This pattern suggests that hippocampal desynchronization during perception may reflect an active reconfiguration of spatial processing rather than a simple reduction in hippocampal engagement. More broadly, these findings align with proposals that low-frequency desynchronization regulates transitions between internally oriented mnemonic states and externally driven perceptual processing [64, 65].

Hippocampal fast theta in humans peaks near 8 Hz, placing it at the boundary of canonical theta and alpha bands, a frequency range where visual neocortex exhibits retinotopic organization in the form of contralateral alpha desynchronization accompanied by HFA increases [43]. The suppression of fast theta by visual stimulation parallels contralateral alpha desynchronization in visual cortex [43], which reflects release of inhibitory gating during active visual processing [66]. Alpha oscillations in visual and temporal regions also track spatial position across saccades and are dynamically coupled to hippocampal rhythms during visual exploration [66, 67]. The theta suppression we observe may thus reflect an analogous release of inhibition in hippocampus. Notably, we did not observe any corresponding HFA increases in a targeted analysis. HFA is thought to index local spiking and mirrors the spatial structure of alpha desynchronization [43]. Thus, the hippocampal visual response appears to be carried entirely by oscillatory suppression without corresponding increases in local activity. This pattern is consistent with fMRI evidence that hippocampus and other default network regions exhibit BOLD decreases during visual mapping [25], and invasive recordings in nonhuman primates confirm that this default network deactivation is retinotopically specific and tracks multiunit activity [68]. This supports the interpretation that the spatially selective theta suppression we observed reflects genuine visuospatial organization in hippocampal circuitry.

The stronger contralateral bias in the right than left hippocampus align with a well-established pattern of right-lateralized spatial processing in the MTL. Neuroimaging studies have demonstrated preferential right MTL and hippocampal engagement during encoding of nonverbal, spatially organized information relative to left MTL engagement for verbal material [69]. Convergent evidence comes from experiments on the effects of brain lesions. Right temporal lobectomies involving large hippocampal excisions specifically impair recall of spatial location on object-location association tasks, an effect not observed following left-sided resections [70]. Moreover, right hippocampal damage selectively impairs visual maze learning [71]. This spatial lateralization extends beyond the visual domain to tactile tasks [72, 73], indicating that right hippocampal specialization for spatial processing is a general property of its circuitry rather than a modality-specific adaptation. The predominance of slow theta and its stronger contralateral bias in the right hemisphere in our experiment suggests that this lateralization has a specific oscillatory signature, and that the neural basis of right hippocampal spatial specialization partly reflects a hemispheric asymmetry in the distribution and spatial sensitivity of slow theta oscillations. More broadly, this finding implies that laterality in hippocampal function is not reducible to the verbal/nonverbal distinction but extends to the spatial precision of theta.

We considered fixational eye movements as potential confounds. Although theta modulations were not predicted by microsaccade rate, spontaneous microsaccades can reflect covert shifts of spatial attention that occur independently of overt visual input [74]. This is relevant because hippocampal theta power is sensitive to attentional state and stimulus expectancy [44, 45], and a spurious association between spontaneous microsaccades and contralateral spatial attention could in principle mimic a visual field bias. The absence of any relationship between microsaccade rate and either slow or fast theta argues against this account and strengthens the interpretation that the contralateral bias we observed reflects visual field sensitivity.

Several limitations of the current study merit consideration. Our sample comprised patients with medically intractable epilepsy. Abnormal hippocampal tissue near seizure onset zones may alter oscillatory properties or their visual sensitivity. Because participants maintained central fixation from a fixed position, the spatial reference frame underlying the observed visual field biases cannot be unambiguously determined. The observed contralateral hemifield bias is equally consistent with head-centered or other egocentric reference frames known to organize the firing of hippocampal neurons [32, 75]. Distinguishing between these possibilities would require paradigms that dissociate these reference frames. Finally, the pRF models we used were translated from fMRI-based retinotopic mapping designed for use in early and intermediate cortex, and thus are not optimized for oscillatory responses in hippocampus. However, the range of model fits are consistent with fMRI-based pRF studies of high-order regions such as the default network [25] and hippocampus [23], which have been interpreted as evidence for visual field biases rather than retinotopic maps [17].

The present findings reframe how hippocampal theta should be understood in relation to the visual system. Rather than treating theta as a signal primarily indexing navigational computation [1, 2] or memory processing [4– 6], our results show that theta carries spatially organized visual input, without any overt navigational or mnemonic demand. This visual sensitivity may provide the spatial scaffold upon which hippocampal circuits construct scene memories, cognitive maps, and object-location associations–the functions with which hippocampal theta has long been associated [76]. More broadly, by demonstrating that hippocampal theta is organized by the spatial structure of visual input in a manner analogous to higher visual cortex, our findings situate the hippocampus firmly within the visual processing hierarchy and provide an electrophysiological foundation for the synthesis of perceptual and mnemonic accounts of human hippocampal function.

## Methods

### Participants

Twelve patients (7 females; aged 45 *±* 16 years (mean *±* SD), range 22-70 years) undergoing presurgical monitoring for intractable epilepsy via iEEG were recruited. All participants had stereotactic depth electrodes implanted in the hippocampus to determine seizure onset zones. Additional monitoring outside of the hippocampus was performed using subdural strips and/or intracranial depth electrodes. Data were collected at The University of Chicago Medical Center (Chicago, IL) and Northwestern Memorial Hospital (Chicago, IL). Research protocols were approved by the Institutional Review Board at both sites prior to data collection. Participants provided written informed consent to take part in the study.

### Experimental paradigm

Six retinotopic mapping runs were collected per participant similar to those performed by Benson et al. [20]. During each run, wedge, bar, or ring apertures slowly moved across the visual field, revealing dynamic backgrounds containing colored objects from different categories overlaid on a 1/f pink noise background. These stimuli were chosen because they elicit responses within the inferior temporal and early visual cortices [20, 61, 62], as well as the hippocampus [23, 26]. During retinotopic mapping runs, participants fixated on a central semitransparent dot that changed colors every 1-5 seconds. Participants were tasked with indicating via key press whenever there was a color change. Images of example stimulus displays presented in Figures 1, 3, and 4 are taken from [20] and are publicly available (https://kendrickkay.net/analyzePRF/, https://www.cns.nyu.edu/kianilab/Datasets.html).

### Intracranial EEG recordings

Stereotactic EEG electrodes (contacts spaced 5-10 mm apart, AD-TECH Medical Instrument Co., Racine WI and DIXI Medical, Marchaux-Chaudefontaine, France) were implanted for clinical seizure monitoring. Signals were recorded using a Neuralynx ATLAS recording system (Bozeman, MT), Nihon Kohden amplifier, or a Natus Quantum system, at a sampling rate of at least 2 kHz, referenced during acquisition to a scalp contact. Data were offline re-referenced to a bipolar montage and downsampled to 500 Hz prior to analysis. Zero-phase IIR notch filters removed 60 Hz line noise and its harmonics up to the Nyquist frequency.

To exclude epileptiform activity, interictal discharges (IEDs) were detected as large transient increases in signal envelope amplitude [77], and data epochs within 1,000 ms of any IED were discarded. Channels with greater than 60% of data near IEDs were excluded from analysis, resulting in 9 of 108 hippocampal contacts removed. Of the remaining data, 21% of epochs were excluded due to IEDs.

### Eye movement recordings

Eye movements were recorded with an EyeLink 1000 Plus remote tracking system (SR Research, Ontario, Canada) at either 500 Hz (N=2 participants) or 1000 Hz (N=10 participants), depending on the recording setup at each site. Before each run, gaze was calibrated and validated using a five-point grid. Calibration was performed until eye tracking validation error was within 1° of visual angle (0.60° *±* 0.34°, mean *±* SD). Continuous gaze records were parsed into fixation, saccade, and blink events using motion (0.15°), velocity (30°/s), and acceleration (8000°/s^2^) thresholds. Blinks were identified based on pupil size, and remaining epochs below detection thresholds were classified as fixations.

Microsaccades were identified from periods of fixation identified by the EyeLink parser, as specified above. For each fixation, we estimated the standard deviation of the horizontal and vertical velocities of the gaze position using a median-based estimator [54, 55]. We applied a velocity threshold of six times the standard deviation, averaging the threshold across temporally neighboring fixations to reduce noise in the estimate that may occur for fixations of shorter duration. Within each fixation, microsaccades were identified as consecutive samples exceeding the velocity threshold for at least 6 ms. The rate of microsaccades was computed as the number of detected microsaccade events per the total duration of fixation periods.

### Electrode localization

Electrode locations within the hippocampus were confirmed with visual inspection, following our prior work [49, 67, 78]. Post-implant computed tomography (CT) images were coregistered to pre-surgical T1-weighted structural MRIs using SPM12 [79]. T1-weighted MRI scans were normalized to MNI (Montreal Neurological Institute) space by using a combination of affine and nonlinear registration steps, bias correction, and segmentation into gray matter, white matter, and cerebrospinal fluid components [80]. Deformations from the normalization procedure were applied to individual electrode locations identified on post-implant CT images using Bioimage Suite [81].

### Separation of oscillatory and broadband neural activity

To identify the neural processes underlying potential retinotopic organization in the hippocampus, we fitted power spectra observed during the task into a combination of aperiodic and periodic components [82]. Estimates of spectral power were obtained through Morlet wavelet convolution (six cycles) at 80 frequencies from 1 to 200 Hz. We adapted a robust regression procedure [49, 82] to fit the log of power and frequency across both the entire task blocks and shorter events of interest within the retinotopy task. Aperiodic 1/f signal fitting was performed for power computed at frequencies between 1.5 Hz to 45 Hz. Aperiodic 1/f power was subtracted from their respective frequencies in the total power spectrum to provide whitened power spectra. After fitting the aperiodic 1/f signal, we analyzed activity within slow (1.5 Hz - 5.0 Hz) and fast (6.0 Hz - 12.0 Hz) theta bands using power from the whitened spectra. To quantify broadband high-frequency activity (HFA, frequencies between 80 Hz - 200 Hz), power values were first log_10_ transformed, normalized across time, then averaged across frequencies. These power values do not come from the whitened power spectra.

To characterize spatial patterns of low-frequency oscillations across hippocampal contacts, power spectra (1 to 40 Hz) were decomposed using non-negative matrix factorization (NMF [83, 84]). Prior to decomposition, the aperiodic component (fitted as described above) was removed from each contact’s spectrum, residuals were baseline shifted to ensure non-negativity, and *L*_2_ normalization was applied across frequencies. The number of NMF factors (*k*) was selected via speckled cross-validation [85], wherein entries of the data matrix were randomly held out (30% per fold, 10 folds), iteratively imputed from the current low-rank estimate, and the held-out mean squared error was evaluated for possible values of *k* from 2 to 10. The lowest value of *k* that fell within at least one standard error above the minimum was selected (*k* = 8, see Fig. S1A). The final factorization was obtained using an alternating least-squares algorithm (*n* = 1, 000 replications). Stability of the NMF solution was assessed using a consensus co-assignment matrix (Fig. S1F) with an overall cophenetic correlation coefficient of 0.87.

After using NMF to identify oscillatory factors, we partitioned recording contacts into clusters based on the NMF weight space (W) with k-means clustering using the squared Euclidean distance. The number of clusters (*k*) was determined by silhouette analysis evaluating *k* from 2 to 8. Stability of the solution was quantified with the bootstrapped adjusted rand index [86] (*n* = 1, 000 interactions of contact-level resampling with replacement).

### pRF modeling

pRFs were estimated using a three stage coarse-to-fine fitting procedure with the CSS model [58]. The CSS model predicts neural responses to the visual stimuli by computing the gain-scaled dot product between a stimulus aperture and a 2D isotropic Gaussian, then applying a compressive static power-law nonlinearity (n *<* 1). A baseline intercept was included to account for general differences in activity in the absence of visual stimulation across contacts. Unlike fMRI applications of this model, no hemodynamic response function was applied. Models were fit separately to predict changes in slow and fast theta power. The fitting procedure proceeded in three stages: first, x-position, y-position, gain, and pRF size were estimated; second, these fits seeded an additional fit for the baseline intercept; third, fits from the second stage seeded a final fit for the compressive exponent. Model performance was evaluated using in-sample r-squared.

### Statistical analysis

#### Stimulus-dependence of microsaccades

To assess whether microsaccades were preferentially directed toward the stimulus aperture, we computed the cosine of the angle between each microsaccade vector (defined by the start and end locations of the microsaccade) and the vector from the start of the microsaccade to the aperture centroid. Values near 1 indicate movements toward the aperture, and values near -1 indicate movements away. We summarized directionality per participant as the mean cosine across all microsaccades, and tested whether this value exceeded chance using a permutation approach (5,000 iterations, shuffling microsaccade directions within each participant). We applied the same test to axis alignment (|cos*θ*|).

#### Anatomical organization of theta oscillations

Hemispheric lateralization and long-axis differences in cluster membership were tested against within-subject spatial permutation null distributions (10, 000 permutations), with all *p*-values FDR-corrected within each family of tests. For the long-axis comparison, we used the Spearman rank correlation between each contact’s MNI Y-coordinate and its binary cluster membership (slow vs. fast theta) as the test statistic. With two clusters this is equivalent to a Mann-Whitney U test [87] of anterior-posterior position between groups. Permuting coordinates within each patient controlled for individual differences in electrode coverage.

#### Effects of visual stimulation on hippocampal activity

To minimize autocorrelation between task epochs, data were divided into events defined as the minimum number of frames in which a given pixel on the screen was continuously stimulated (50 frames at 30 Hz, approximately 1667 ms). We first tested the effects of stimulus presence, stimulus coverage, and stimulus laterality on theta oscillations at the individual contact level. To do this, we performed multivariate nonparametric permutation testing (10, 000 permutations) using the Freedman-Lane procedure [88]. First, we fitted a full general linear model (GLM) to estimate the observed t-statistic for each variable of interest. Next, for each variable, we fitted a reduced GLM excluding the variable of interest while retaining remaining variables as nuisance regressors, and computed the residuals from this reduced model. These residuals thus reflect variance unexplained by the remaining nuisance regressors. For each permutation, the residuals from the reduced model were randomly permuted, and added back fitted values of the reduced model, and the full GLM was refit to the permuted data. The resulting t-statistic for each regression coefficient formed a null distribution against which the observed t-statistics were compared. Significant contacts were identified when the observed t-statistic exceeded 95% of this null distribution.

This approach was used to identify effects of stimulus presence, coverage, and laterality on slow theta, high theta, and HFA at the contact level. We did not find reliable effects of visual stimulation on HFA. As such, we focused our reporting and group-level modeling on slow and fast theta.

To examine stimulus effects at the group level, we fitted a linear mixed effects model separately for slow and fast theta. Random slopes for fixed effects were initially included to allow for within-subject effects to vary across participants. However, models with these terms failed to converge due to complexity of the random effects structure [89, 90]. Following recommendations in dealing with model convergence [91], random slopes were removed as follows:

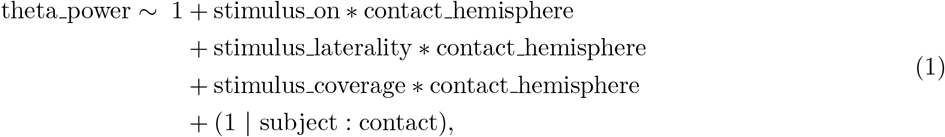

where theta power denotes the average slow or fast theta power within an event; stimulus on indicates whether a stimulus was present; stimulus laterality is the difference in stimulus coverage between ipsilateral and contralateral hemifields relative to the recorded hippocampal contact; stimulus coverage is the total proportion of the visual field occupied by the stimulus (left plus right hemifields combined); and contact hemisphere is the hemisphere of the hippocampal contact. Fixed effects were tests with parametric bootstrap likelihood ratio tests, and p-values were Holm-Bonferroni corrected [92] across the two theta bands.

We used this model and bootstrap approach to simultaneously address three primary questions: whether hippocampal theta tracks stimulus presence, whether it is sensitive to how much of the visual field is stimulated, and whether it reflects the relative balance of ipsilateral versus contralateral stimulation. We also used this model to assess differences between left and right hippocampi.

#### Comparison of slow and fast theta activity

To test whether the stimulus effects described above differed between theta frequencies (see Equation 1), we fitted an extended model with theta frequency (i.e., slow or fast theta) as an interactive factor:

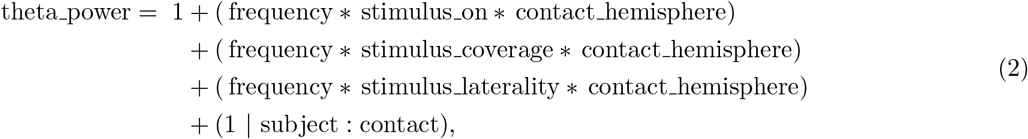

where frequency indicates slow or fast theta, and all other terms are defined as in Equation 1. Statistical inference followed the procedures as defined above.

#### Behavioral influences on theta activity

We tested whether ongoing behavior during the task (button presses and microsaccades) modulated hippocampal theta. Data were divided into windows between task-related color changes. We labeled each window according to the presence or absence of a button press indicating a color change, and the presence or absence of microsaccades. Then we fitted the following model to predict changes in whitened theta power:

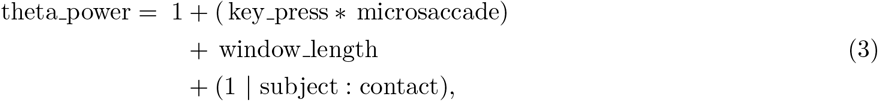

where key press and microsaccade are categorical factors indicating occurrence of an event, and window length is an additional covariate to account for variable durations between color changes. As with prior models, fixed effects were tested with parametric bootstrap likelihood ratio tests and Holm-Bonferroni corrected across theta bands.

## Supplementary Figures

**Figure S1:**
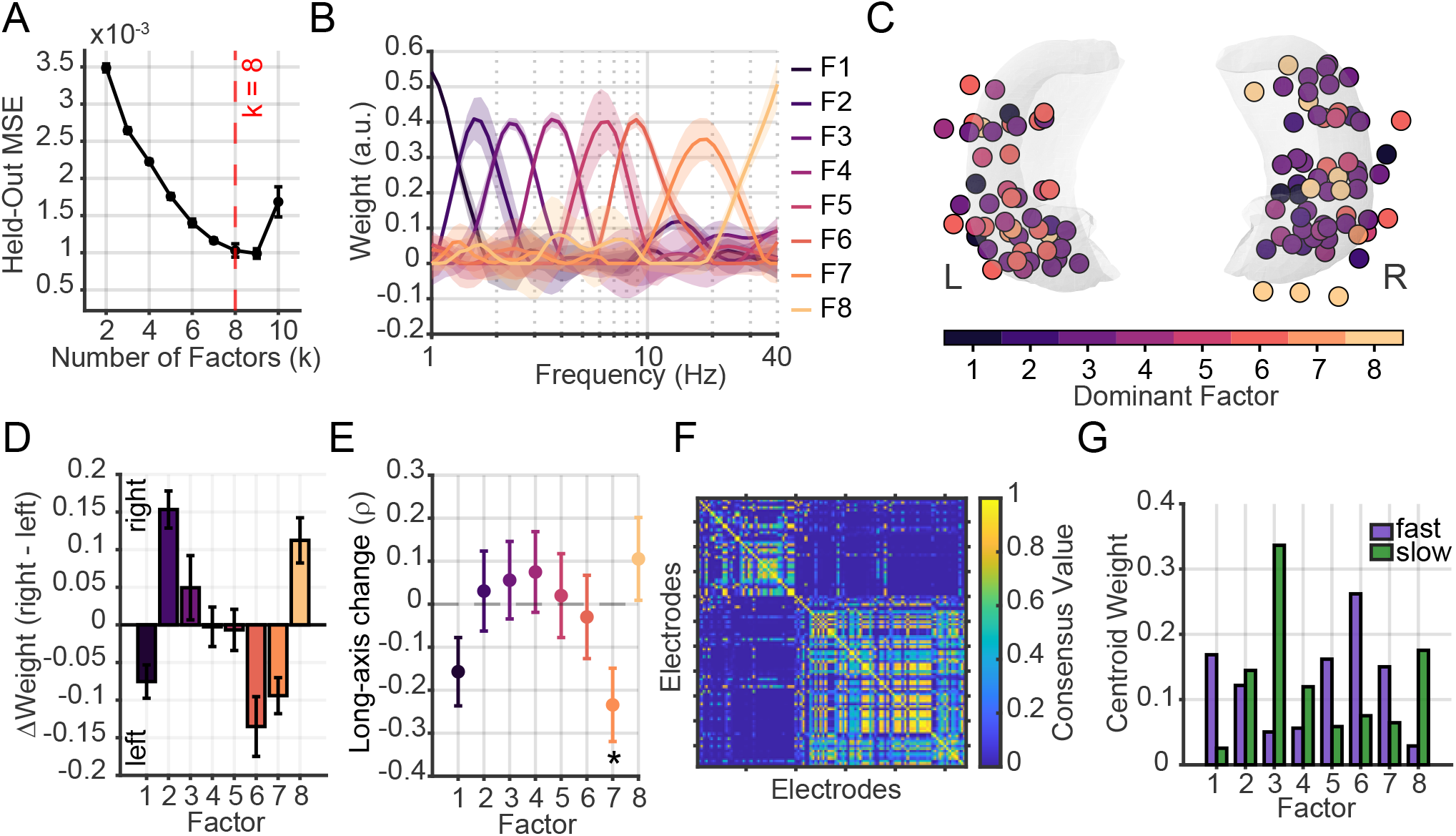
Characterization of hippocampal oscillations using non-negative matrix factorization (NMF) and k-means clustering. (**A**) Selection of k=8 factors, based on held-out mean squared error (MSE) from 10-fold cross validation. Error bars denote *±*1 standard error across folds. The dashed red line marks the smallest k whose MSE lies within one standard error of the minimum. (**B**) Average spectral profiles (H matrix) for each of the eight factors across frequencies (1–40 Hz). Shaded bands denote bootstrap-estimated SEM across subjects, illustrating the reproducibility of each factor’s peak frequency. (**C**) Winner-takes-all spatial map showing, for each of the *N* = 108 contacts, the factor with the greatest loading weight. Electrodes are displayed on a hippocampal surface for the left (L) and right (R) hemispheres; color denotes dominant factor. (**D**) Hemisphere asymmetry of factor weights, expressed as the difference in mean loading between right and left hippocampus (ΔWeight = right − left). Error bars denote *±*1 SEM estimated by within-subject resampling. No factor showed a significant hemispheric difference (all *p >* 0.05, FDR-corrected). (**E**) Long-axis gradient for each factor, quantified as the Spearman correlation (*ρ*) between factor loading weight and anterior-posterior coordinate in MNI space. Points show observed *ρ*; error bars denote *±*1 SEM estimated by within-subject resampling. Only factor 7 (peak ∼18 Hz, beta band) showed a significant long-axis gradient, with greater loadings in anterior hippocampus (*ρ* = −0.23*±*0.09, *p* = 0.037, FDR-corrected, ∗). (**F**) Consensus co-assignment matrix. Each entry reflects the proportion of runs in which two electrodes received the same winner-takes-all cluster assignment; values range from 0 (never co-clustered; dark blue) to 1 (always co-clustered; yellow). Rows and columns are ordered by average-linkage hierarchical clustering of the 1− consensus distance matrix. The strong block-diagonal structure indicates highly reproducible cluster separation across initializations. (**G**) Normalized centroid weight profiles for the fast theta (green) and slow theta (purple) clusters. Each bar shows the mean factor loading weight at the cluster centroid, normalized to sum to one. The slow theta cluster is dominated by factor 3 (peak ∼2.5 Hz) and the fast theta cluster by factor 6 (peak ∼8.8 Hz).

